# Machine Learning Uncovers the Transcriptional Regulatory Network for the Production Host *Streptomyces albidoflavus*

**DOI:** 10.1101/2024.01.09.574332

**Authors:** Mathias Jönsson, Renata Sigrist, Tetiana Gren, Mykhaylo Semenov Petrov, Nils Marcussen, Anna Svetlova, Pep Charusanti, Peter Gockel, Bernhard O. Palsson, Lei Yang, Emre Özdemir

## Abstract

*Streptomyces albidoflavus* is a popular and genetically tractable platform strain used for natural product discovery and production via the expression of heterologous biosynthetic gene clusters (BGCs). However, its transcriptional regulatory network (TRN) and its impact on secondary metabolism is poorly understood. Here we characterized its TRN by applying an independent component analysis to a compendium of 218 high quality RNA-seq transcriptomes from both in-house and public sources spanning 88 unique growth conditions. We obtained 78 independently modulated sets of genes (iModulons) that quantitatively describe the TRN and its activity state across diverse conditions. Through analyses of condition-dependent TRN activity states, we (i) describe how the TRN adapts to different growth conditions, (ii) conduct a cross-species iModulon comparison, uncovering shared features and unique characteristics of the TRN across lineages, (iii) detail the transcriptional activation of several endogenous BGCs, including surugamide, minimycin and paulomycin, and (iv) infer potential functions of 40% of the uncharacterized genes in the *S. albidoflavus* genome. Our findings provide a comprehensive and quantitative understanding of the TRN of *S. albidoflavus*, providing a knowledge base for further exploration and experimental validation.

**Graphical Abstract:** 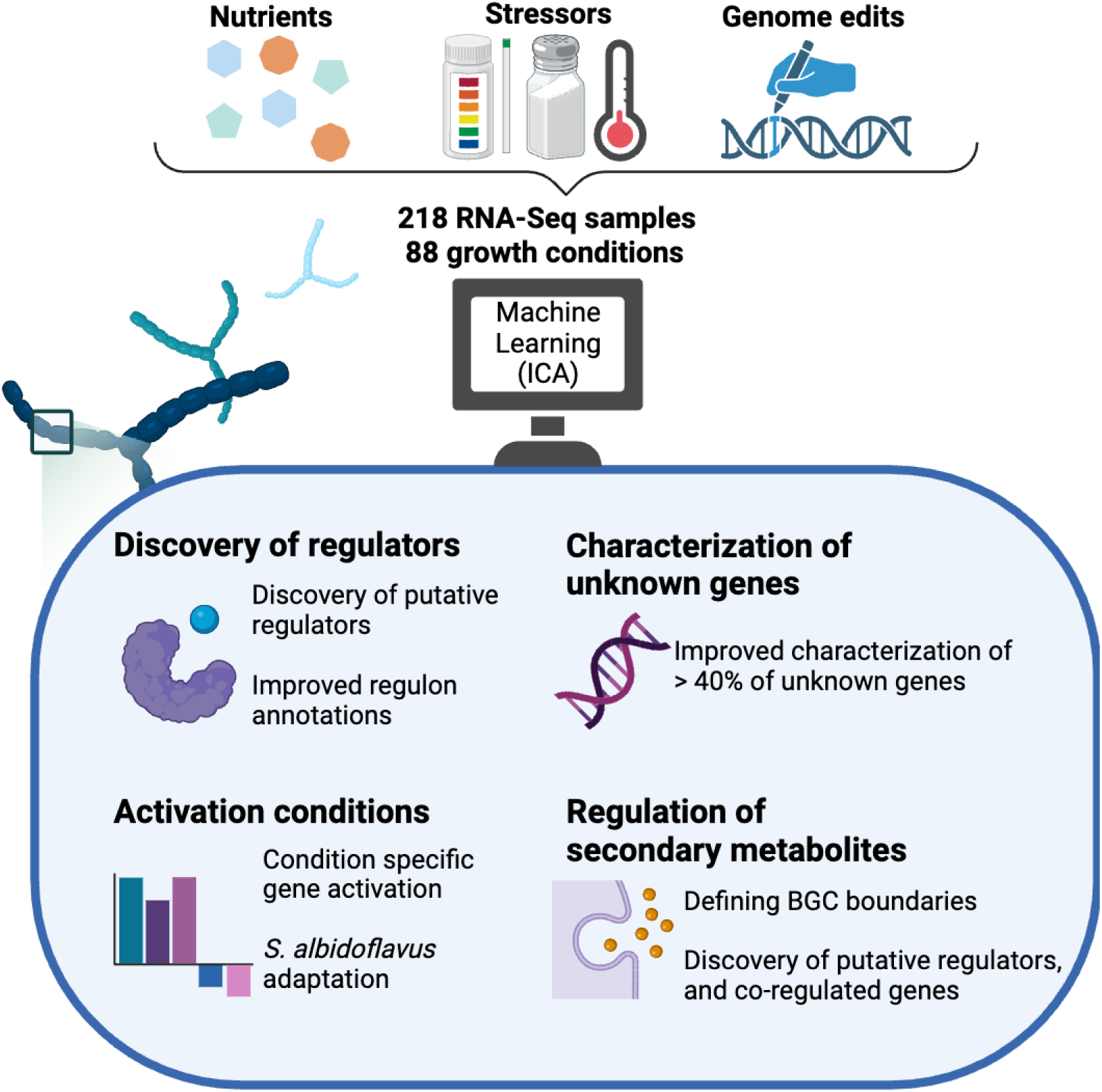

## Introduction

*Streptomyces* is a well-known genus of Gram-positive bacteria capable of producing diverse secondary (specialized) metabolites, including antibiotics, anticancer agents, and antifungals (1). These products are encoded in biosynthetic gene clusters (BGCs), with most *Streptomyces* species encoding 20-30 BGCs (2–5); Many BGCs, however, are not expressed in laboratory settings. Consequently, a more thorough understanding of the regulation of these BGCs may lead to activation of more BGCs and the discovery of more novel compounds (6, 7).

The production of secondary metabolites is controlled, in part, by the transcriptional regulatory network (TRN), which controls the expression of genes in response to genetic and environmental changes. *Streptomyces* genomes are often large (6 - 12 Mbp), and bottom-up approaches to characterize the TRN have revealed an intricate hierarchy of regulators — ranging from global to pleiotropic and local — that activate under specific growth conditions and developmental stages, reflecting the complex life cycle of *Streptomyces* (8–13). *Streptomyces albidoflavus* J1074 (previously *Streptomyces albus* J1074) is a well characterized and widely used heterologous host for studying cryptic Actinomycete BGCs and producing industrially important secondary metabolites (14–23). *S. albidoflavus* has one of the smallest *Streptomyces* genomes of 6.8 Mbp, containing around 23 endogenous BGCs (24).

Moreover, it contains a large proportion of genomic regulators, including over 400 transcription factors and around 30 sigma factors (24). The number of potential regulators together with the large genome present significant challenges for bottom-up approaches in characterizing the TRN on a large-scale.

Independent Components Analysis (ICA) is a powerful unsupervised machine learning method that provides a top-down approach to characterizing the TRN (25, 26). It decomposes gene expression data into sets of independently modulated sets of genes (iModulons) and their corresponding activity profiles across different conditions (27–29). ICA has been used to infer genome-scale TRNs and functional modules for several bacterial species, including *Escherichia coli*, *Bacillus subtilis*, and *Mycobacterium tuberculosis,* often with strong overlap with regulons detailed in the literature (28–32). Importantly, the modularization of the transcriptome into independent signals has enabled the iModulons to be comparable and transferable across species, enabling new pathways in host species, showcasing great potential for engineering microbial cell factories and biomanufacturing (33).

To gain a comprehensive and systematic understanding of the TRN of *S. albidoflavus*, we performed ICA on a compendium of 218 high-quality gene expression profiles, of which 161 were newly generated for this purpose. We identified 78 iModulons which reveal novel insights into the transcriptional responses of *S. albidoflavus* to relevant conditions, such as nutrient limitation, BGC removals, heterologous expression, and to various stressors. We also conduct a large-scale comparison of iModulon structure across seven different organisms to validate our findings and to gain an evolutionary context of the TRN. Our analysis reveals novel insights into BGC regulation, identifying co-regulated genes, defining BGC borders, and putative regulators. To promote the use of this knowledge base, all iModulons can be searched, browsed, and studied through imodulondb.org.

## Methods

### RNA extraction and library preparation

We assessed *S. albidoflavus* activity across various environmental and nutritional factors using Biolog microplates. Based on the metabolic activities observed, we identified the most promising conditions that supported robust growth and activated diverse metabolic pathways (Supplementary Table 1.1). Next, we imitated these favorable conditions during our *S. albidoflavus* culturing process for subsequent RNA extraction. We grew *S. albidoflavus* and extracted RNA from 211 samples, each with at least one biological replicate, under 83 unique growth conditions. These conditions included supplemented minimal media with different carbon sources, salinity, pH, and temperature stresses, genome-reduced strain (up to 10 endogenous BGCs deleted), and heterologous expression of actinorhodin BGC. Genome-reduced strains were obtained by deleting 10 BGCs from *S. albidoflavus* parental strain using CRISPR-Cas9 (34) or CASCADE-Cas3 systems (35). Additionally, actinorhodin BGC was integrated via conjugation into the chromosome of parental and genome reduced hosts.

*S. albidoflavus* strains were cultivated aerobically in ISP2 (4 g/L yeast extract, 10 g/L malt extract, 4 g/L dextrose, and distilled water, pH 7), DNPM (40 g/L dextrin, 7.5 g/L bacteriological peptone from meat, 5 g/L yeast extract, 21 g/L MOPS, 750 mL miliQ water, 250 mL tap water, pH 7) or minimal medium (6 g/L (NH_4_)_2_SO_4_, 1.86 g/L citric acid, 0.21 g/L FeCl_3_, 3.33 mL/L 2M MgSO_4_, 250 g/L of monosodium glutamate, 0.1 M phosphate buffer pH 7.0 and 1.6 mL/L trace metal solution (36) (20.4 g/L H_2_SO_4_ 96%, 50 g/L citrate·H_2_O, 16.75 g/L ZnSO_4_·7H_2_O, 2.5 g/L CuSO_4_·5H_2_O, 1.5 g/L MnCl_2_·4H_2_O, 2 g/L H_3_BO_3_, and 2 g/L NaMoO_4_·2H_2_O). The recipe for DNPM typically contains soytone but was replaced with bacteriological peptone from meat in our formulation (15). In conditions where cultures were grown on solid media, plates containing Mannitol Soya Flour (MS) medium (20g/L Mannitol, 20g/L Soya Flour, 20g/L Agar, and tap water, pH 7) were used. Solid culture samples were recovered using 10 µL inoculation loops. In culture conditions where different carbon sources were tested, monosodium glutamate was replaced by selected carbon sources at 66 mM. In salinity stress conditions, NaCl solutions were used to supplement the correspondent medium. In temperature heat stress conditions, cultures were submerged in a ScanVac HeatSafe 18 water bath set to 42C°. For routine cultivation, the temperature was set up at 30C°.

Total RNA was extracted using the RNeasy Mini QIAcube kit (Qiagen; cat. No./ ID: 74116) and treated with DNase I according to the manufacturer’s instructions. RNA concentration and integrity were verified using Qubit 2.0 (Invitrogen) and Fragment analyzer (Agilent 530V). rRNA purification, library preparation and RNA-Seq were performed by Azenta Life Sciences. NEBNext rRNA Depletion kit (Bacteria) (New England Biolabs; cat. No. E7850X) was used for rRNA depletion. Barcoded libraries were prepared for each condition in duplicate with NEBNext Ultra II Directional RNA Library Prep Kit for Illumina using default protocols. Libraries for all samples were sequenced using Illumina NovaSeq 6000.

### Biolog Phenotype Microarrays

*S. albidoflavus* activity across various nutritional sources was evaluated by Phenotype MicroArray^TM^ analysis. Uniform spore suspension (80% T, transmittance) was obtained from 5-7 days sporulated MS plates and used to prepare inoculating fluids according to Biolog protocol for *Streptomyces.* IF-0a base inoculating fluid and Redox Dye Mix D were used for this experiment. PM 1-2 (carbon sources), 3 (nitrogen sources) and 4 (phosphorus and sulfur sources) Biolog microplates were selected and inoculated with 100 µL of prepared inoculation fluid. PM microplates were incubated in OmniLog® PM System, at 30°C, 72h.

Data analysis was performed with DuctApe (0.18.2) (37) using dphenome module as described on manual pages. In brief, phenotype microarray data for *S. albidoflavus* was loaded, background signals based on control wells were subtracted, signals were trimmed, data was analyzed based on default 10 clusters and inconsistent replicates were purged.

### Data acquisition and processing

Data processing and quality control is detailed in Sastry et al. (38) and available at https://github.com/SBRG/iModulonMiner. Briefly, read trimming was performed using Trim Galore with the default options (https://www.bioinformatics.babraham.ac.uk/projects/trim_galore/), followed by FastQC (https://www.bioinformatics.babraham.ac.uk/projects/fastqc/). Next, reads were aligned to the *S. albidoflavus* reference genome from NCBI (NC_020990.1) using Bowtie (39). The read direction was inferred using RSEQC (40) before generating read counts using featureCounts (41). Lastly, all quality control metrics were compiled using MultiQC (42).

Samples that failed any of the four FASTQC metrics were discarded: per base sequence quality, per sequence quality scores, per base n content, and adapter content. Furthermore, samples with less than 500 000 reads mapped to coding sequences were removed, and hierarchical clustering was used to identify any sample outliers. Only samples with at least two replicates and a Pearson R correlation of > 0.90 between replicates were analyzed. Furthermore, to minimize possible batch effects, each individual experiment was normalized to a reference condition prior to calculating the iModulons (Supplementary Table 1.2) (38). After quality control and normalization, the final data compendium comprised 218 high-quality expression profiles: 161 generated from this study and 57 collected from NCBI-SRA. The two public datasets that met our quality control criteria included PRJNA1003853, which tested the heterologous expression of Nybomycin in a *S. albidoflavus* strain with 14 endogenous BGCs removed (43), and PRJNA708335, which heterologously expressed pamamycin in the wild type *S. albidoflavus* (44).

### Performing Independent component analysis (ICA)

Independent components analysis was implemented as per McConn et al. (45) on the RNA-Seq compendium. Specifically, the scikit-learn implementation of FastICA was executed 100 times with random seeds and a convergence tolerance of 10^−7^ (46, 47). The resulting independent components (ICs) were clustered using DBSCAN to identify robust ICs, using an epsilon of 0.1 and a minimum cluster seed size of 50. To account for identical components with opposite signs, the following distance metric was used for computing the distance matrix:

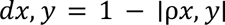

where ρ*x*, *y* is the Pearson correlation between components *x* and *y*. The final robust ICs were defined as the centroids of the cluster.

Since the number of dimensions of ICA can affect the modularization, we applied the above procedure to the dataset multiple times, ranging the number of dimensions from 10 to 190 (i.e. the approximate size of the dataset) with a step size of 10. 190 dimensions generated the most robust iModulons, and were used for this analysis.

### Cross-species comparison of iModulons

To enable comparisons of iModulon structures between organisms we first identified orthologous genes among seven species represented on imodulondb.org, (*Escherichia coli, Salmonella enterica, Pseudomonas aeruginosa, Bacillus subtilis, Mycobacterium tuberculosis, S. albidoflavus*, and *Sulfolobus acidocaldarius*) using Orthofinder with default settings (48). These species were chosen to represent a diverse range of phyla, spanning both closely related and distantly related species. All iModulon data files and objects were downloaded from imodulondb.org (28). The gene weight files of all species describing the iModulon memberships were merged based on orthogroups identified by orthofinder. If more than one gene per species were present in a given orthogroup only the highest gene weight was kept per iModulon. Next, we conducted cosine similarity in a pairwise manner for all iModulons, and visualized the similarities using Cytoscape (49). Edge weights were filtered (0.648) to only show the top 1500 edge weights, and we used the MCL cluster algorithm from the clusterMaker app to identify clusters of iModulons (50).

### Characterization of iModulons

To facilitate iModulon characterization, we used the PyModulon Python package (51). *k*-means clustering was used to identify component-specific thresholds, as per PyModulon recommendation. Manual editing of some iModulon thresholds were done to improve characterization efforts. The iModulons were characterized by information from public databases, literature and based on similarities to other iModulons from other species in our iModulome analysis. Gene annotations and operon information was downloaded from Biocyc.org (52). BGC regions were annotated using antiSMASH with default settings (53). Code for our analysis pipeline and description of iModulons is maintained on github (https://github.com/biosustain/salb_imodulons).

## Results

### Independent component analysis reveals 78 iModulons

To capture a wide range of expression states we cultured *S. albidoflavus* in 68 unique growth conditions. These conditions included different stressors, genome modifications, and nutrient sources, which were chosen based on activity screening using Biolog Phenotype Microarrays (Supplementary Table 1.1). This in-house generated dataset comprises 161 high quality RNA-Seq transcriptomes, increasing publicly available RNA-Seq datasets from SRA NCBI by 190% (Figure 1A and Supplementary Table 1.2). We combined our in-house generated dataset with public datasets from the SRA database, and after quality control, the final compendium consisted of 218 samples from 88 unique growth conditions, with a high median Pearson R score of 0.99 between replicates (Figure 1B, C).

**Figure 1.**
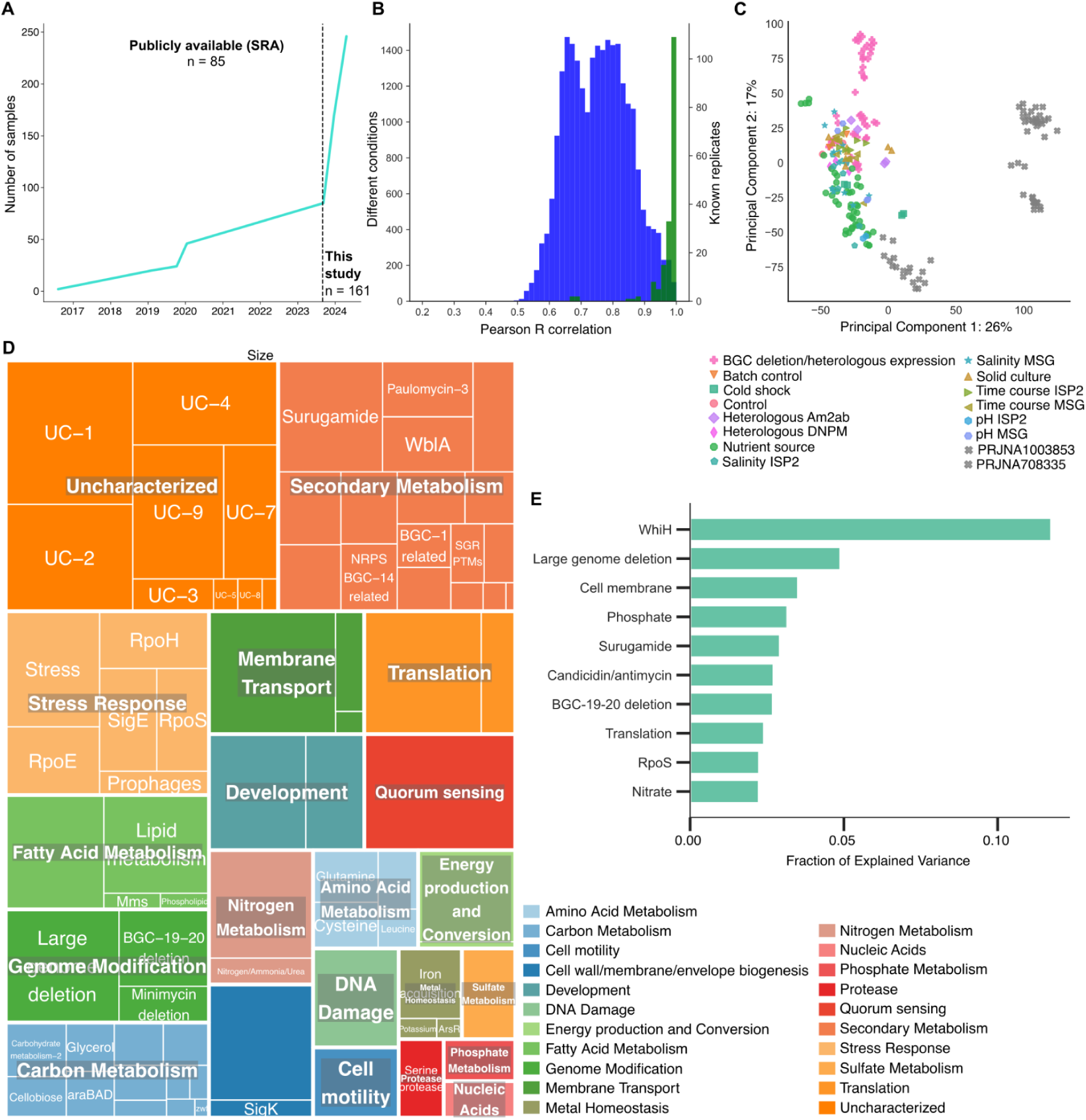
Overview of the S. albidoflavus iModulons. (**A**) The availability of public RNA-Seq (SRA) data for S. albidoflavus over time. (**B**) Histogram of the correlations between log-tpm profiles of all samples across all conditions (blue) and known replicates (green). Median Pearson R score between replicates is 0.99. Replicates that did not exceed 0.90 in R score were removed from further analysis. (**C**) Principal components analyses of 218 samples used for ICA. The color and symbol depicts the specific project where the sample was generated. All in-house generated samples for this study are colored, and the two public datasets included in this study are labeled in gray. (**D**) A treemap depiction of the 78 identified iModulons and their associated functional categories. The dimensions of the individual boxes correlate with the number of genes found in each respective iModulon. (**E**) Highlight of the top 10 iModulons contributing the greatest proportion of explained variance. These iModulons represent key functions such as development, translation, secondary metabolite biosynthesis, and stress.

ICA decomposed the RNA-Seq compendium into 78 iModulons (Figure 1D), where each iModulon contains a set of genes whose expression levels vary concurrently with each other, but independently of all other genes not in the given iModulon.

Overall, 48.4% of all genes were assigned into an iModulon, explaining 79% of the variance in the gene expression (Figure 1E, Supplementary Table 1.3 and Supplementary Table 1.4). The iModulons cover a wide range of functions, including secondary metabolism (21.8%), carbon metabolism (15.4%), stress response (7.7%), as well as, currently uncharacterized sets (11.5%) (Figure 1D). We have provided the information for each iModulon in the form of an interactive dashboard on imodulondb.org (28). The dashboard is a user-friendly way for researchers to search for or browse the details of iModulons or genes of interest.

### Cross-species iModulon comparison help characterize new iModulons

Given the limited number of regulons described for *S. albidoflavus* in the literature, we developed a method to compare and validate its iModulon structures with those of other species available on iModulonDB. We identified orthologous gene weight similarities across all iModulons from seven species, including *S. albidoflavus*, and visualized the iModulons in a similarity based network that cluster together based on their function, regardless of species, which we refer to as the iModulome (Figure 2A, Supplementary Table 2.1-2.5 and Supplementary Figure 1). Of the 78 *S. albidoflavus* iModulons, 94.9% clustered together with at least one other iModulon allowing us to compare and characterize the respective gene contents.

**Figure 2.**
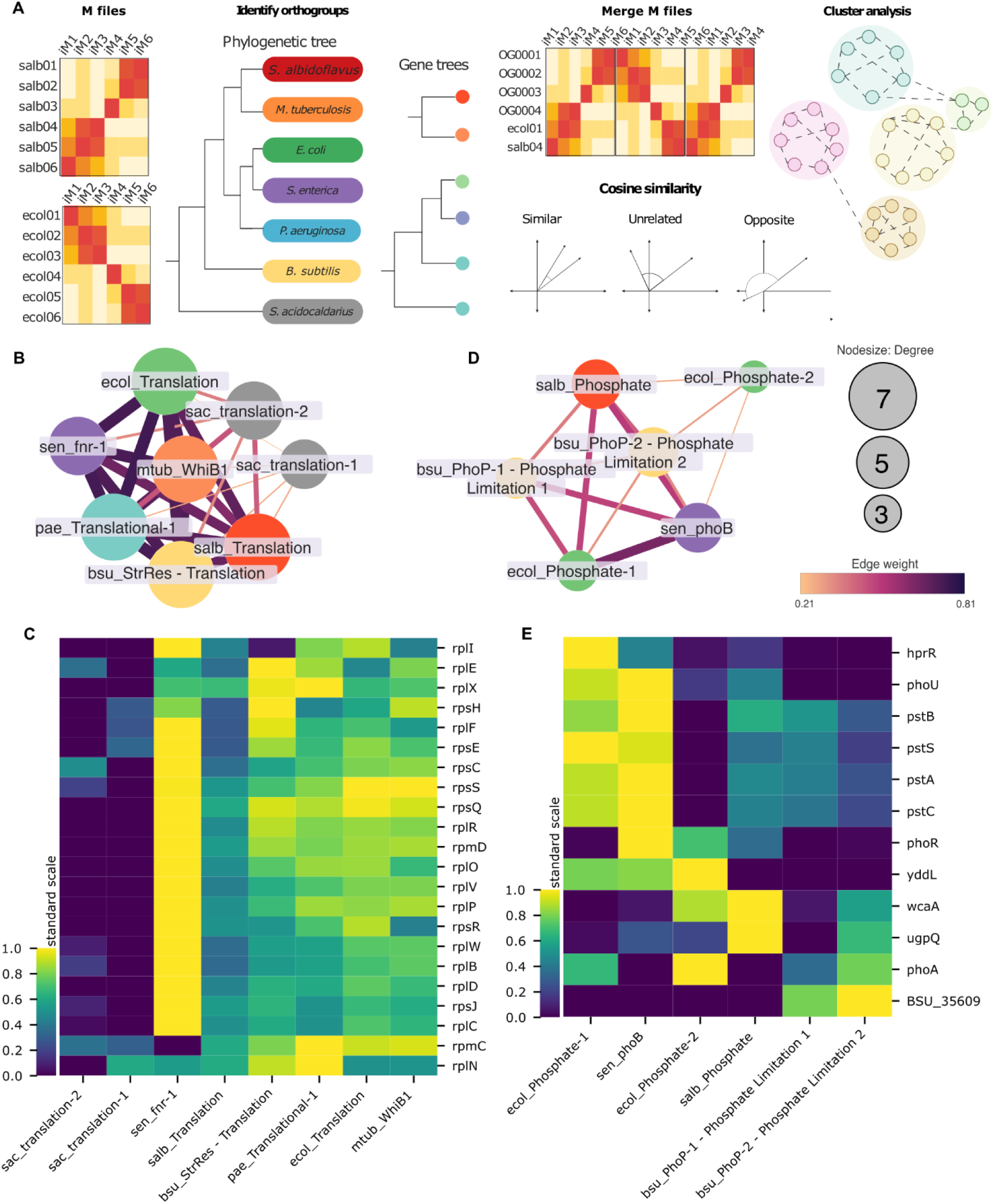
S. albidoflavus iModulons similarities and differences with other organisms. (**A**) Overview of the construction of the iModulome. M files generated from ICA analysis of seven organisms were collected. Orthogroups were identified using orthofinder (48), and used to merge the M files based on the specific orthogroup. Cosine similarity score and MCL clustering was used to create a clustered network of iModulons based on similarity. (**B**) A cluster from the iModulome of iModulons across seven species (coloured nodes; Red, S. albidoflavus; Orange, M. tuberculosis; Green, E. coli; Purple, S. enterica; Blue, P. aeruginosa; Yellow, B. subtilis; Grey, S. acidocaldarius) based on cosine similarity of ICA derived gene weights, containing iModulons related to translation processes. (**C**) Clustermap of core genes in the cluster, the scale bar depicts the standardized weights of orthologous genes, expressed as z-scores (**D**) A cluster of phosphate related iModulons. (**E**) Clustermap of core orthologous genes in the Phosphate related cluster.

For example, the Translation-related cluster of the iModulome comprises eight iModulons from the seven included species, with a high average edge weight of 0.52, indicating significant similarity in the gene weights of orthologous genes (Figure 2B). This high similarity is largely based on the expression patterns of 11 core genes, which all encode for conserved ribosomal subunits (Figure 2C). *S. acidocaldarius* has two iModulons in this cluster, but they are the least connected, with an average edge weight of 0.31 to the other iModulons, indicating their more distant phylogeny. Moreover, the phosphate-related cluster of the iModulome consists of six iModulons from four species, with an average edge weight of 0.36 (Figure 2D). We identified the core genes as related to the *pstSCAB* and *PhoPR* systems, however there is greater heterogeneity regarding co-expressed genes, suggesting potential lineage-specific adaptations to gene expression under phosphate-related conditions (Figure 2E).

### iModulons give insights into the regulation and activation of BGCs

Of the 23 predicted BGC regions in *S. albidoflavus*, 12 had core biosynthetic genes as members of 16 iModulons, including paulomycin, cyclofaulknamycin, candicidin, and surugamide (Table 1). The remaining BGCs had evenly distributed transcriptional activity across all conditions, and thus were not modularized by ICA. The iModulons provide a comprehensive overview of the expression of the core biosynthetic genes across different growth conditions along with co-regulated genes, both within and outside of the BGC. For example, the BGC regions predicted by ICA were 62.7% smaller compared to antiSMASH predictions, with Surugamide F having the lowest coverage (4.55%) and Streptamidine covering 100% of the antiSMASH predicted region (Supplementary Figure 2). Notable co-regulated genes located outside of the respective BGCs included putative regulators, amino acid transporters, and cytochrome P450 enzymes (Supplementary Table 1.4). Furthermore, we detected enriched DNA binding motifs that appear upstream of the operons in BGC-related iModulons, further suggesting co-regulation (Table 1 and https://github.com/biosustain/salb_imodulons). Below we analyze some BGC-related iModulons in more detail, to provide novel insights into their potential regulatory mechanisms and expression.

**Table 1.**
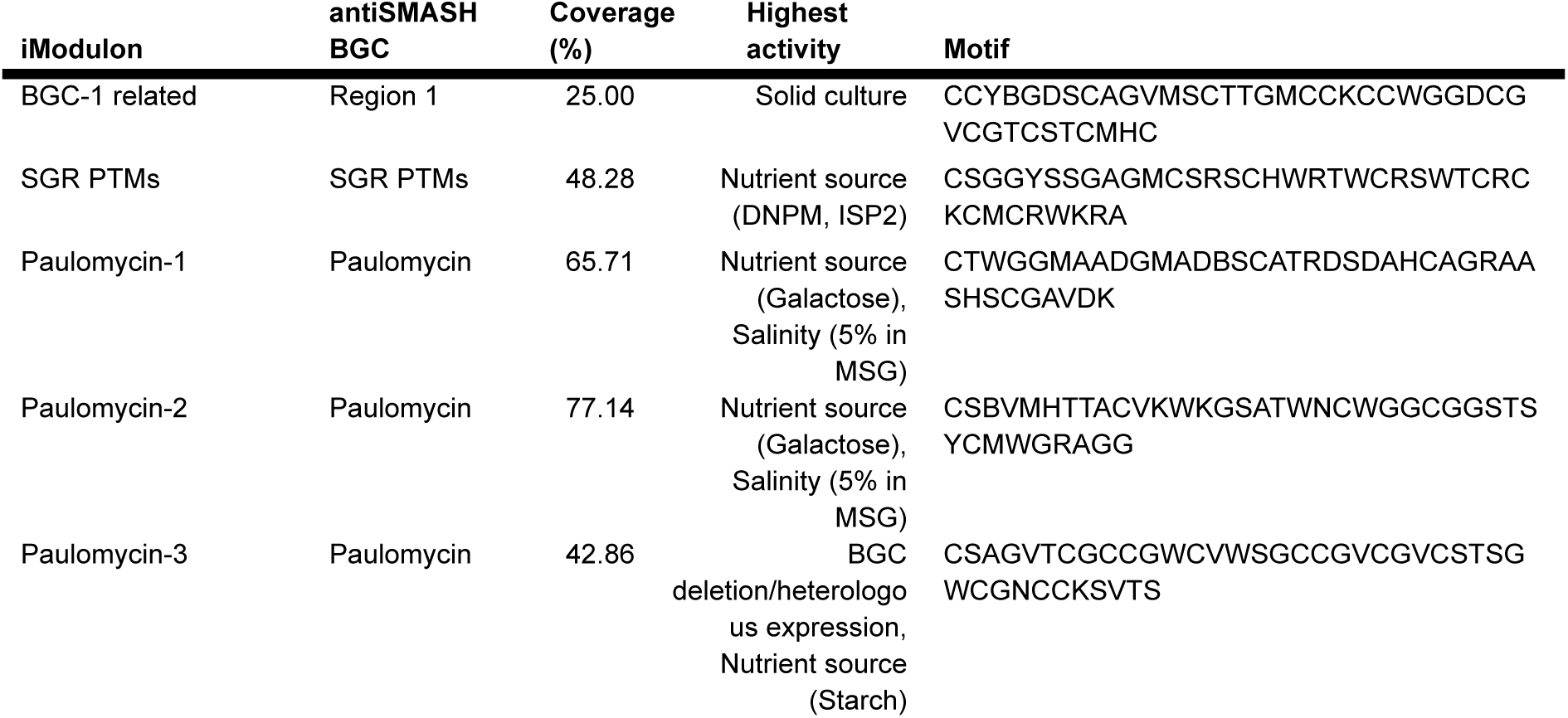
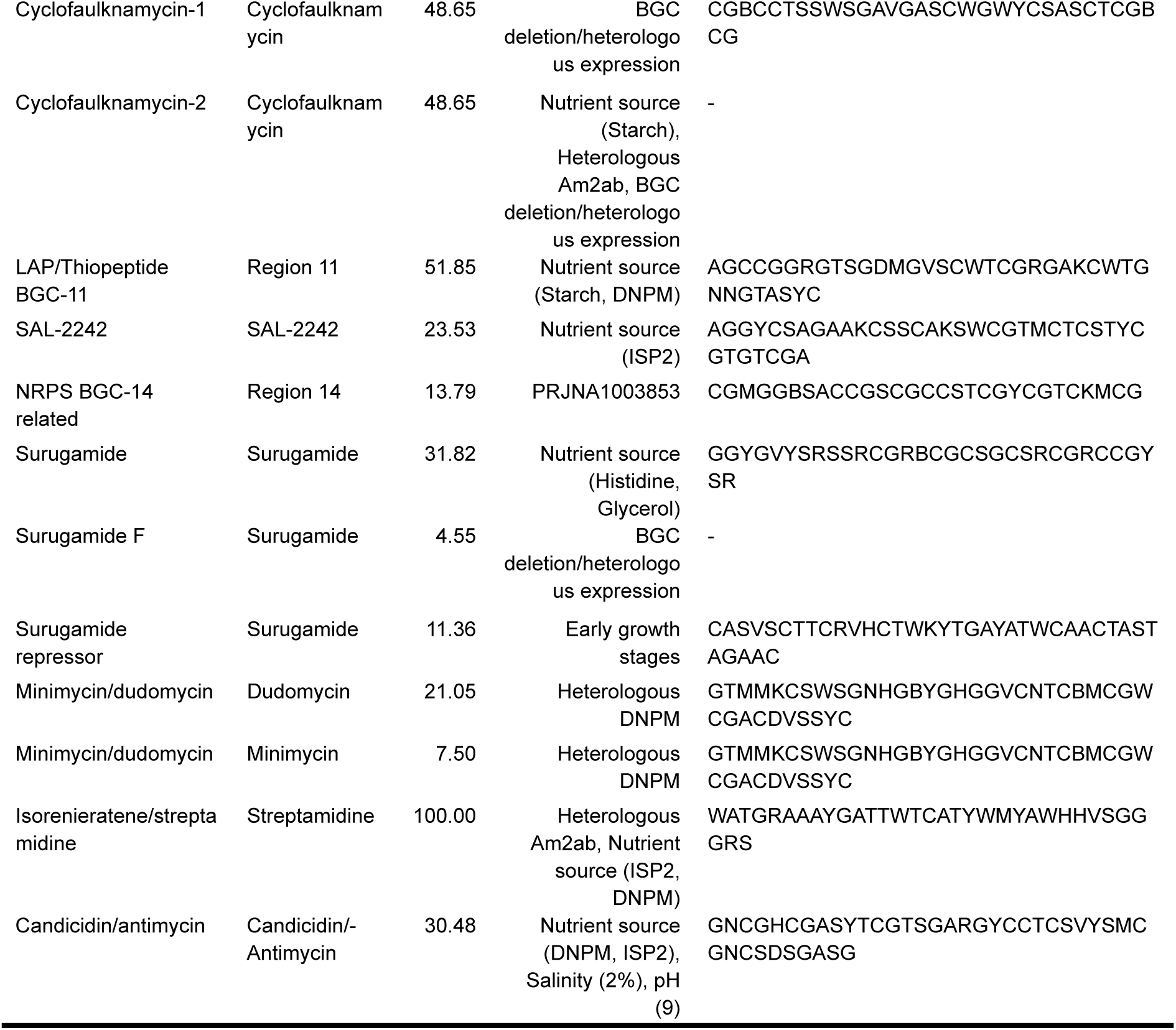
Biosynthetic gene cluster representation in iModulons. List of antiSMASH predicted BGC regions and corresponding iModulons. The coverage represents the percentage of genes from the antiSMASH predicted BGC region that is present in the iModulon. The highest activity column, depicts the growth conditions of the samples with the highest iModulon activation scores. The motif column shows the IUPAC ambiguity codes of the top enriched motif for the operons and genes in the iModulon.

### Novel insights into the regulation of surugamide

We identified three iModulons that contain core biosynthetic genes for surugamide, a cyclic non-ribosomal peptide synthetase (NRPS) with antibacterial and antifungal properties (54, 55). These iModulons, named Surugamide, Surugamide Repressor, and Surugamide F, consist of 89, 7, and 2 genes, respectively (Figure 3A, B).

**Figure 3.**
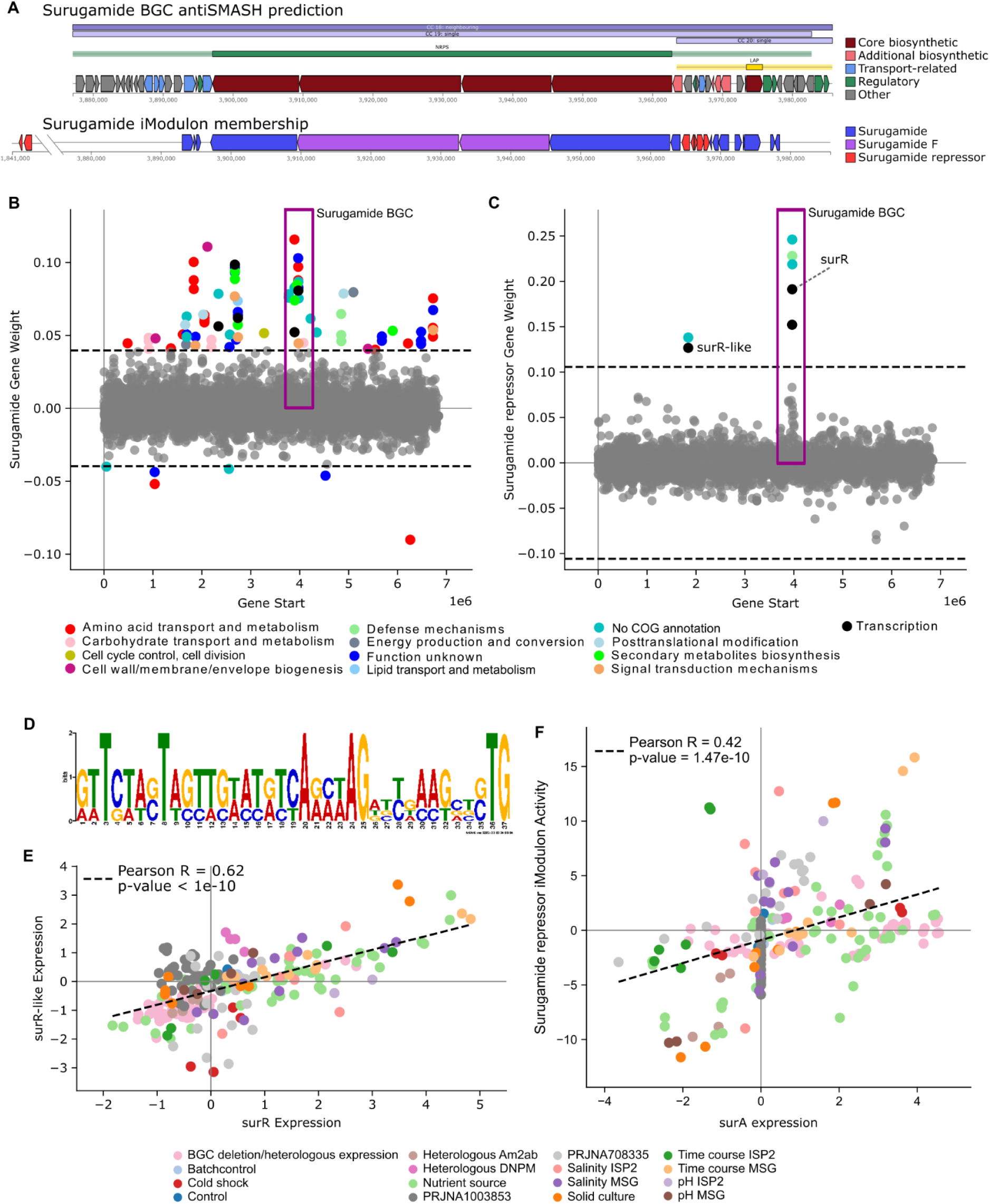
Surugamide activation and repression. (**A**) Gene locations of the surugamide BGC region as depicted by antiSMASH v7.10.0 and the gene memberships in corresponding iModulons. (**B**) Gene weights of the Surugamide iModulon. The y-axis depicts the individual gene weightings, and x-axis shows the location of the gene in the genome. Genes within the surugamide BGC are located within the purplebox. Colour depicts COG category. 22.5% of the genes in this iModulon are related to amino acid transport and metabolism. (**C**) Gene weights for the Surugamide-repressor iModulon, which contains the regulator surR and a surR-like located 2Mbp downstream of the surugamide BGC. (**D**) Sequence logo of a significantly enriched DNA binding motif in the Surugamide-repressor iModulon. (**E**) Scatterplot depicting the correlation in gene expression across all samples for surR and the surR-like, indicating similar expression patterns across most growth conditions. (**F**) Scatterplot illustrating the correlation between the iModulon activity of Surugamide repressor and the gene expression of surA.

Surugamide F appears to be an artifact due to partial BGC deletion in certain conditions (Supplementary Figure 3). The Surugamide and Surugamide F iModulons comprise 13 and 2 genes, respectively, from the predicted BGC region, including the core biosynthetic genes *surABCD*, transporters, and putative regulators (Figure 3A). Additionally, 22.5% of the Surugamide iModulon genes are related to amino acid transport and metabolism, along with four co-regulated cytochrome P450 genes.

In contrast, the Surugamide Repressor iModulon contains the previously characterized repressor *surR*, which regulates transcription of the *surABCD* operon (Figure 3B) (56). In addition, we identified an undescribed *surR*-like gene located approximately 2 Mbp downstream of the surugamide BGC (Figure 3C). Both *surR* and the *surR*-like factor exhibit similar expression patterns across various conditions and we identified an enriched DNA binding motif within this iModulon, suggesting co-regulation (Figure 3D and 3E). Surprisingly, neither *surR* or *surR*-like are significantly correlated with any genes from the *surABCD* operon, suggesting that there are likely other factors involved in the regulation of the surugamide BGC (Figure 3F).

### Identification of co-regulated gene clusters with two NRPS BGCs

The Minimycin/dudomycin iModulon contains core biosynthetic genes for two NRPS BGCs predicted by antiSMASH, minimycin (also known as oxazinomycin) and a dudomycin-like (Figure 4A and 4B). This iModulon is downregulated under salinity stress in minimal media (Figure 4C). In addition to the two BGCs, this iModulon contains two additional gene clusters, one that appears to be involved in amino acid transport, with genes such as *ilvE*, aminotransferase, and tryptophan 2,3-dioxygenases, and one that contains genes such as *ectB*, SidA/IucD/PvdA family monooxygenase, and an AMP-binding enzyme, suggesting these gene clusters may be important for the biosynthesis and transport of necessary components of the biosynthesis of minimycin and the dudomycin-like.

**Figure 4.**
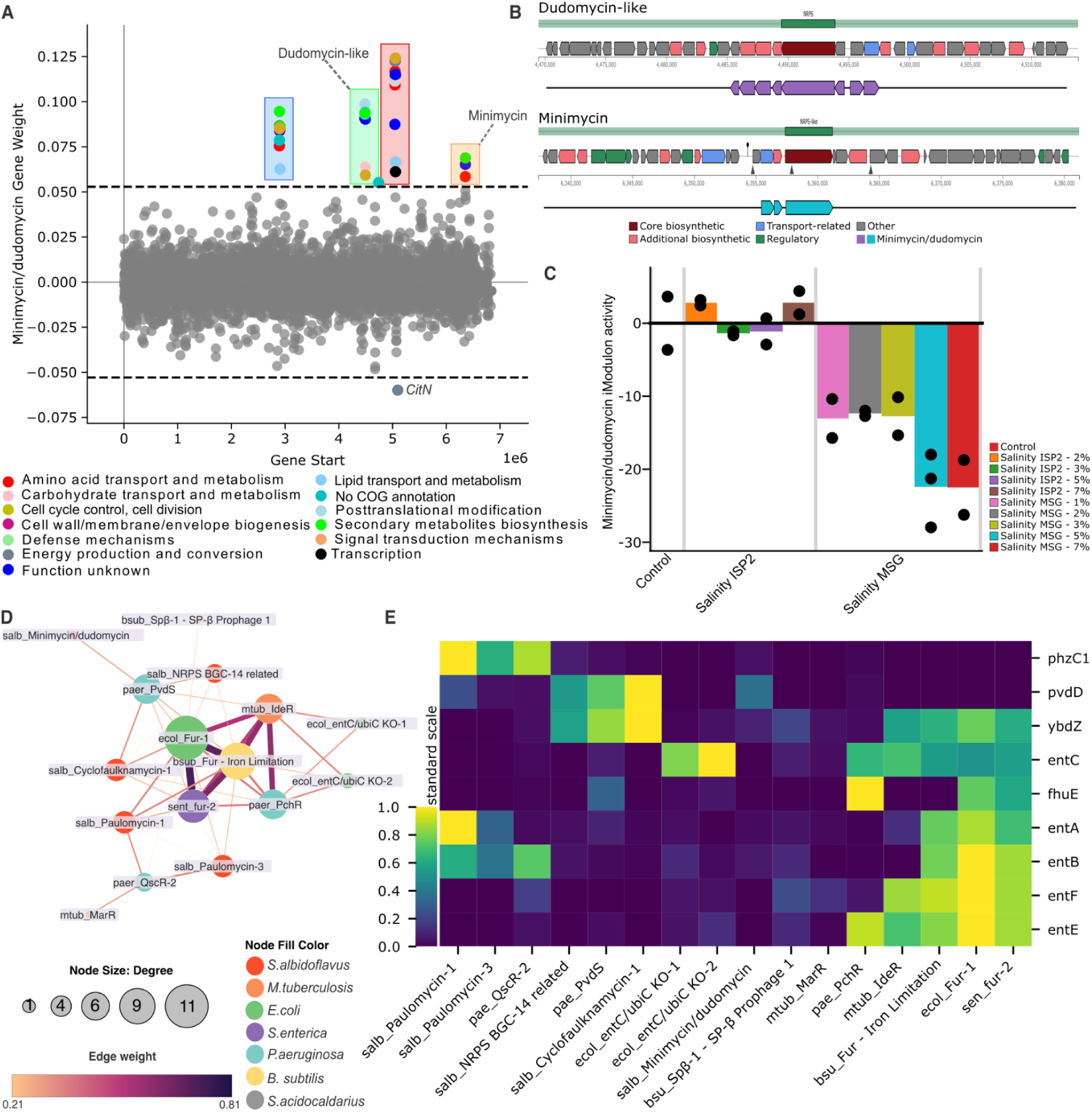
NRPS activation in iron limited conditions. (**A**) Gene weights of the Minimycin/dudomycin iModulon, where two distinct gene clusters, in addition to the two BGCs, are highlighted. Genes are plotted against their position in the genome. The color of the dots depicts the COG function according to the legend. (**B**) Genome location and orientation of the minimycin and dudomycn-like BGCs predicted by antiSMASH, and their coverage within the iModulon. (**C**) Activity plot of the Minimycin/dudomycin iModulon across control and salinity stress conditions. (**D**) A cluster from the iModulome with iModulons related to iron acquisition across many organisms including Five iModulons from S. albidoflavus related to NRPS or PKS-like BGCs. (**E**) Clustermap of core genes in the cluster, which are mainly orthologous genes of the enterobactin synthesis pathway in E. coli.

iModulons involving the NRPS BGCs minimycin/dudomycin, cyclofaulknamycin-1 and NRPS BGC-14 and two iModulons related to the transcription of paulomycin, a polyketide synthase (PKS)-like BGC, cluster with iron limitation-related iModulons of other species in the iModulome (Figure 4D). Core genes of this cluster of iModulons are orthologous of *Fur* regulated and iron dependent genes in *E. coli* involved in the enterobactin biosynthesis pathway, such as *entA*, *entB*, *entC*, *entE*, *entF*, *ybdZ*, and *fhuE*. Additional core genes include orthologs to *phzC1* and *pvdD,* which encode for the phenazine biosynthesis protein *phzC*, and pyoverdine synthetase D, respectively, in *P. aeruginosa* (Figure 4E).

### Uncharacterized iModulons provide a road map to discover novel functions

When analyzing an iModulon covering the BGC for paulomycin, we identified a cluster of four genes located 1.8 Mbp downstream of the paulomycin BGC which is down-regulated in the Paulomycin-1 iModulon (Figure 5A). Interestingly, this cluster consists of one putative GntR family transcription factor and three uncharacterized genes, which together make up a separate uncharacterized iModulon called UC-6 (Figure 5A). Upregulation of UC-6 appears to lower the expression of the Paulomycin BGC in a condition-dependent manner.or instance, cultures grown on solid media appear to upregulate UC-6 while down-regulating Paulomycin-1 (Figure 5B). Characterizing the function of UC-6 and fully understanding its effect on the paulomycin BGC require experimental verification of the uncharacterized genes. The genome of *S. albidoflavus* consists of around 30% with no COG annotation, and many more with no experimental validation (24). Combined, 40% of uncharacterized genes in *S. albidoflavus* were present in at least one iModulon, providing a pathway to validate gene function in future studies (Supplementary Table 1.4).

**Figure 5.**
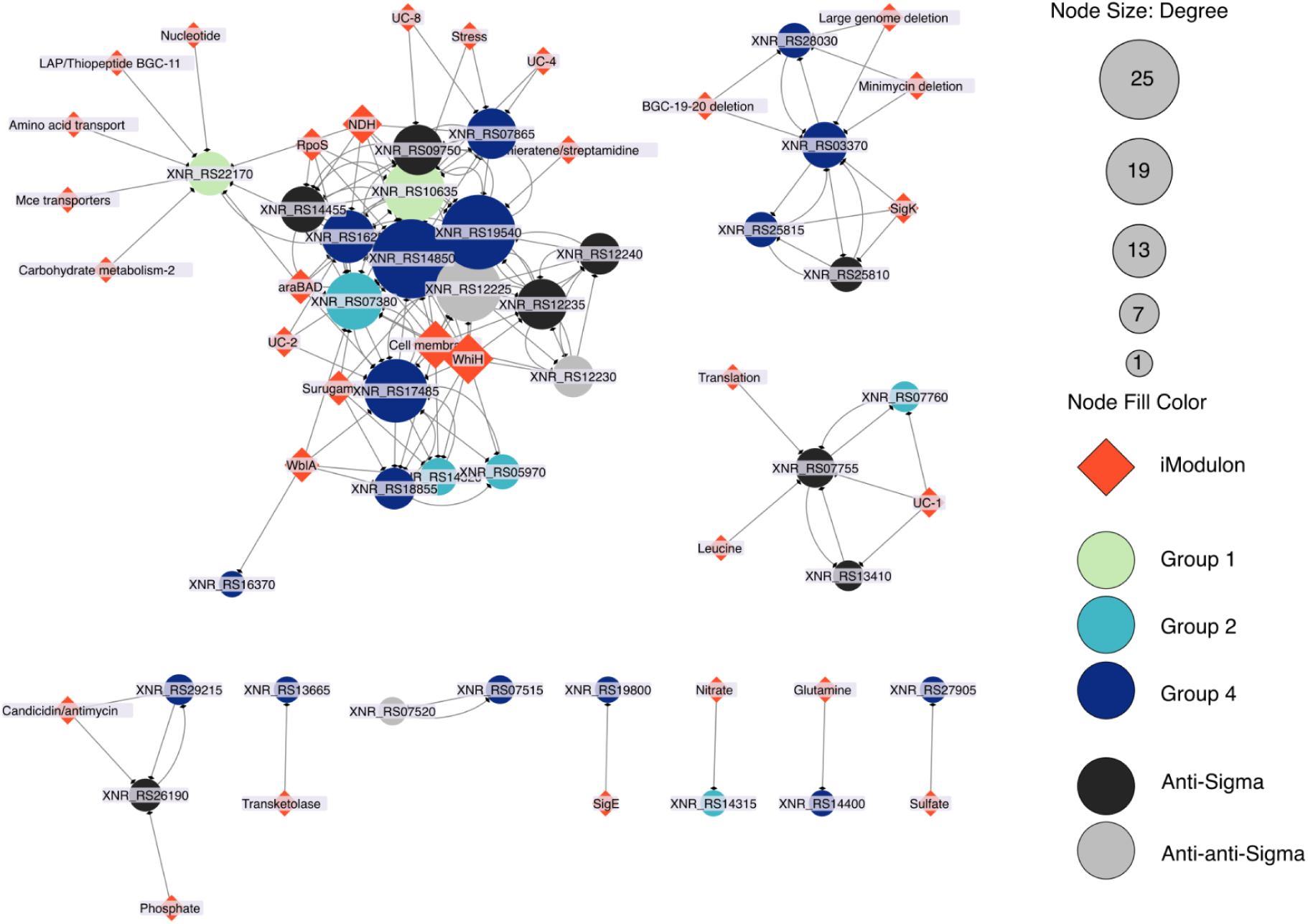
Sigma and related factors activity with iModulons. Gene/iModulon Network analysis of Sigma, anti-sigma, anti-anti-sigma factors (circles), and iModulon (diamonds) activities across the transcriptional compendium. Node size depicts the number of connections in the network. Nodes in the clusters are significantly correlated.

**Figure 6.**
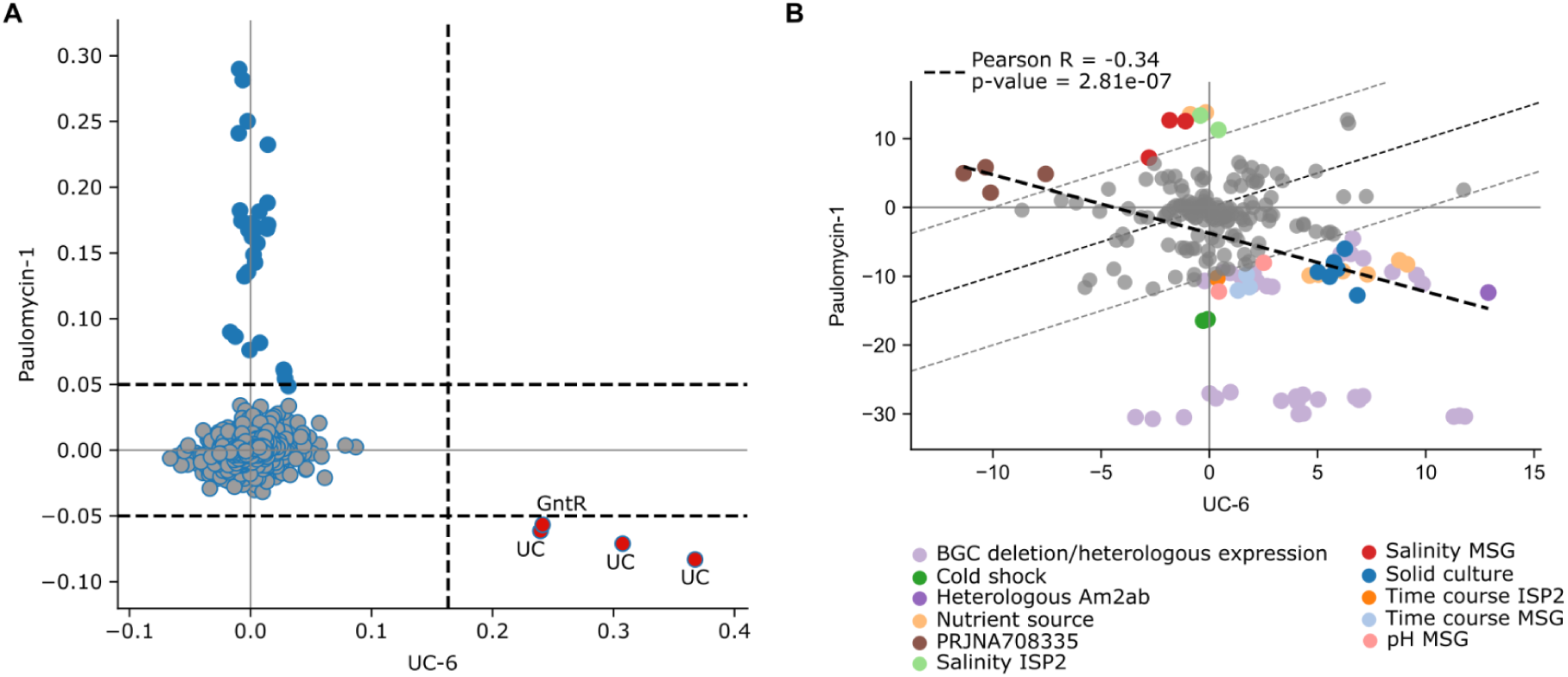
Activities of uncharacterized iModulons. (**A**) Scatterplot of gene weights between the UC-6 and Paulomycin-1 iModulons. Dashed lines indicate the upper and lower thresholds of the respective iModulon. Red genes are members of both iModulons, with high activity in UC-6 and low activity in Paulomycin-1. (**B**) Activity of UC-6 and Paulomycin-1 iModulons across all samples shows conditions which activate the iModulons. Paulomycin-1 is for example down-regulated in solid cultures while UC-6 is upregulated.

### Uncovering the Sigma factor regulatory network of *S. albidoflavus*

Sigma factors are important regulators of gene expression; however, their regulatory roles in *S. albidoflavus* are not well understood, with many of their full regulons not yet being defined (57). We identified 31 sigma, 11 anti-sigma, and three anti-anti-sigma factors annotated in the *S. albidoflavus* genome, with 17 being members of least one iModulon, providing potential regulatory targets (Supplementary Table 3.1). To expand on this, we performed a network analysis by comparing sigma factor activities with iModulon activities across our RNA-Seq compendium (28, 58). This similarity based network revealed 34 sigma-related factors with high expression similarities between themselves or iModulons (Figure 5 and Supplementary Table 3.2). Notably, the expression levels of Group 1 primary Sigma factors *hrdD* (XNR_RS10635) and *hrdA* (XNR_RS22170) exhibited significant correlations with multiple iModulons, including Cell membrane, RpoS, Amino acid transport, and Carbohydrate metabolism-2 (Figure 5). Furthermore, some Group 4 sigma factors such as XNR_RS17485, XNR_RS14850 (*sigT*), and XNR_RS19540, are significantly correlated with several large iModulons, including Surugamide, WblA, WhiH, and RpoS, suggesting potential global regulatory roles for these factors, meriting further investigation.

## Discussion

By applying a machine learning algorithm, ICA, to a large compendium of both in-house generated and publicly available RNA-Seq datasets, we have identified 78 iModulons which provide a systems-level view of the transcriptome and a window into the complex TRN of *S. albidoflavus* (3, 24, 59, 60). Similar to previous studies, this modularization of the TRN resulted in new discoveries, including co-regulation of peripheral gene clusters with BGCs, putative regulators, numerous uncharacterized genes which are promising targets for future research, and provided a detailed compendium of growth conditions and their effect on the TRN (28–32, 58, 61–63). iModulons have started to become valuable tools in strain design and bioproduction, suggesting our analysis will further improve the engineering efforts of *S. albidoflavus* (33). Importantly, the iModulon knowledge base can continuously be updated as new RNA-Seq data becomes available, ensuring more detailed reconstructions of the TRN over time (29, 30, 64). The iModulons are freely available at imodulondb.org.

### The iModulome places the iModulons in an evolutionary context

While many of the *S. albidoflavus* iModulons appear to have significant overlap with known regulons, there remains a comparatively low amount of studies detailing regulators in *S. albidoflavus*, hindering characterization and annotation efforts (10–12, 60). To assist with characterization and to validate the iModulons we introduced the ‘iModulome’ — a comprehensive network of iModulons from seven distinct species, which clusters iModulons together based on orthologous gene weights, allowing a cross-species comparison of gene contents across iModulons. This approach not only facilitated the annotation of iModulons in *S. albidoflavus* but also placed the iModulons in an evolutionary context, opening up a potentially new “pan-modulome” area of research akin to pan-genomics and pan-transcriptomics, which may reveal conserved and lineage-specific aspects of the TRN (65, 66). While this level of analysis was beyond the scope of this study, the similarity between the iModulons of *S. albidoflavus* and those of other species suggests that our results will be broadly applicable to other *Streptomyces* and *Actinomycetes* more generally. Moreover, these methodologies hold promise for application to other non-model organisms to increase our understanding of microbial regulation (62, 67).

### iModulons give insights into the regulation of secondary metabolites

The iModulons captured over half of the predicted endogenous BGCs in the *S. albidoflavus* genome, including polyketide synthases, NRPSs, lanthipeptides, and ribosomally synthesized and post-translationally modified peptide products (RiPPs), detailing growth conditions that transcriptionally up- or down-regulate these clusters. In addition to improving BGC border definitions, the iModulons revealed several co-regulated genes and gene clusters outside the predicted BGC region. Our findings include a *surR*-like transcription factor that is co-expressed with a major regulator of surugamide biosynthesis, *surR,* across the majority of growth conditions (55). In addition, the core biosynthetic genes of surugamide (*surABCD*) are members of an iModulon that include several genes with amino acid transport functions, cytochrome P450 enzymes, and an acyl-CoA dehydrogenase, which is crucial for avenolide activity, a key secondary metabolite signaling molecule (68, 69). Taken together, this iModulon may contain several genes necessary for surugamide biosynthesis, providing a roadmap for experimental validation. Moreover, we identified two off-site gene clusters related to the transcription of the minimycin and dudomycin-like BGCs in *S. albidoflavus*, one which appears involved in amino acid transport and one which contains genes such as *ectB*, SidA/IucD/PvdA family monooxygenase, and an AMP-binding enzyme, suggesting these gene clusters may be important for the biosynthesis and transport of necessary components of the biosynthesis (70, 71). These peripheral sets of genes are likely important when considering strain engineering or BGC engraftments to other organsms, and are promising targets for experimental validation (33). Furthermore, we found that many of the BGC related iModulons were enriched in DNA binding motifs within 500 bp upstream of the BGC, which could be lucrative targets to discover new promoters to increase production yields (72).

Moreover, through the iModulome we identified several BGC-related iModulons that clustered together with *Fur* regulated and iron scavenging-related iModulons of other species (73–75). The *S. albidoflavus* BGCs included cyclofaulknamycin, minimycin and dudomycin-like, an unknown NRPS-like, and the PKS-like paulomycin. While the biosynthesis of minimycin appears to involve a non-heme iron dependent monooxygenase, there is very little known about the involvement of iron in the expression of these BGCs, or potential iron scavenging properties (76–80). This underscores the potential of iModulons as a framework for unraveling the intricate regulatory mechanisms that govern BGC expression, particularly in relation to metal ion homeostasis. The association of these iModulons with iron-dependent pathways not only hints at a broader regulatory landscape but also paves the way for targeted investigations into the iron-mediated control of secondary metabolism.

### iModulons help infer potential functions of uncharacterized genes

Several iModulons were difficult to characterize due to a high number of uncharacterized genes and scarcely available information about regulons. The *S. albidoflavus* genome consists of 30% genes with uncharacterized or unknown functions, and many more that lack experimental validation (24). iModulons have been used to characterize several uncharacterized genes in *E. coli* and have identified many lucrative targets for other species (33, 81). Our analyses enable us to infer potential functions for 40% of all genes that lack a COG annotation, providing promising experimental targets going forward. For instance, several of the iModulons related to BGCs contain co-regulated genes with unknown functions, many of which are located outside of the main BGC, and therefore may be necessary components for biosynthesis. The iModulons generated in this study provides a roadmap to experimentally validate these uncharacterized genes.

### Sigma factors may regulate several iModulons

By analyzing the correlation of activity between iModulons and Sigma factors we generated a network of sigma factors that are potentially orchestrating the regulation of specific iModulons within the *S. albidoflavus* genome. Out of 31 sigma, 11 anti-sigma, and three anti-anti-sigma factors, 17 are integral components of at least one iModulon, and 23 appear significantly correlated with iModulon activities, highlighting the intricate regulatory mechanisms governing gene expression in this species. In addition to the two primary Sigma factors *hrdA* and *hrdD* which appear highly connected and correlated with many iModulons, several extracytoplasmic function sigma factors stood out, including a paralog to *sigT* in *S. coelicolor*, which regulates development and actinorhodin production in response to nitrogen stress (82, 83), and XNR_RS17485 and XNR_RS19540 which are significantly correlated with iModulons related to BGCs, development, and uncharacterized functions. Furthermore, the discovery of enriched DNA binding motifs upstream of each operon within the iModulons provides a valuable blueprint for future research. Studies employing methodologies such as ChIP-, RIViT-, or DAP-seq could leverage our findings to elucidate the direct interactions between these sigma factors and their target genes (84–86). Such investigations would not only validate the regulatory roles suggested by our network analysis but also enhance our understanding of the dynamic control of BGCs, potentially unlocking new avenues for the optimization of secondary metabolite production in *Streptomyces*.

## Conclusion

This is the first study to compile a large compendium of in-house generated and publicly available transcriptomes to identify iModulons in *Streptomyces*, providing a comprehensive and quantitative understanding of the TRN of *S. albidoflavus*. We showed how the iModulons can be used to (i) compare the TRN of *S. albidoflavus* with other organisms on imodulondb, revealing both conserved and lineage-specific features, (ii) elucidate the regulation of BGCs, such as surugamide, minimycin, and paulomycin by identifying putative regulators, co-regulated genes, and condition-dependent activation patterns, and (iii) infer the functions of 40% of the uncharacterized genes in the genome, by associating them with iModulons. Our findings pave the way for future studies to extend and improve the iModulon framework by incorporating new conditions and targeted mutants for transcription factors. Moreover, our results suggest new strategies for rational strain and process design to optimize the production of valuable natural products by *S. albidoflavus* and other *Streptomyces* species.

## Data Availability

All RNA-seq raw files generated from this study are available from NCBI at bioproject PRJNA1062162. Source codes for iModulon analysis and figures are available at https://github.com/biosustain/salb_imodulons.

## Author Contributions

MJ performed computational analysis, generated presented results, and wrote the manuscript. RS, TG, MSP, NM, AS, PC, PG developed protocols and performed experiments. BP, LY, EO conceptualized the study. All authors contributed to interpretation of results, reviewed and/or edited the manuscript.

## Acknowledgements

We are grateful to Tilmann Weber, Kai Blin, Thomas Booth and Simon Shaw for the informative discussions and input when characterizing the iModulons, and to Edward Catoiu for integrating the dataset to imodulondb.

