## Supplementary figure 1 for "Machine Learning Uncovers the Transcriptional Regulatory Network for the Production Host *Streptomyces albidoflavus*"

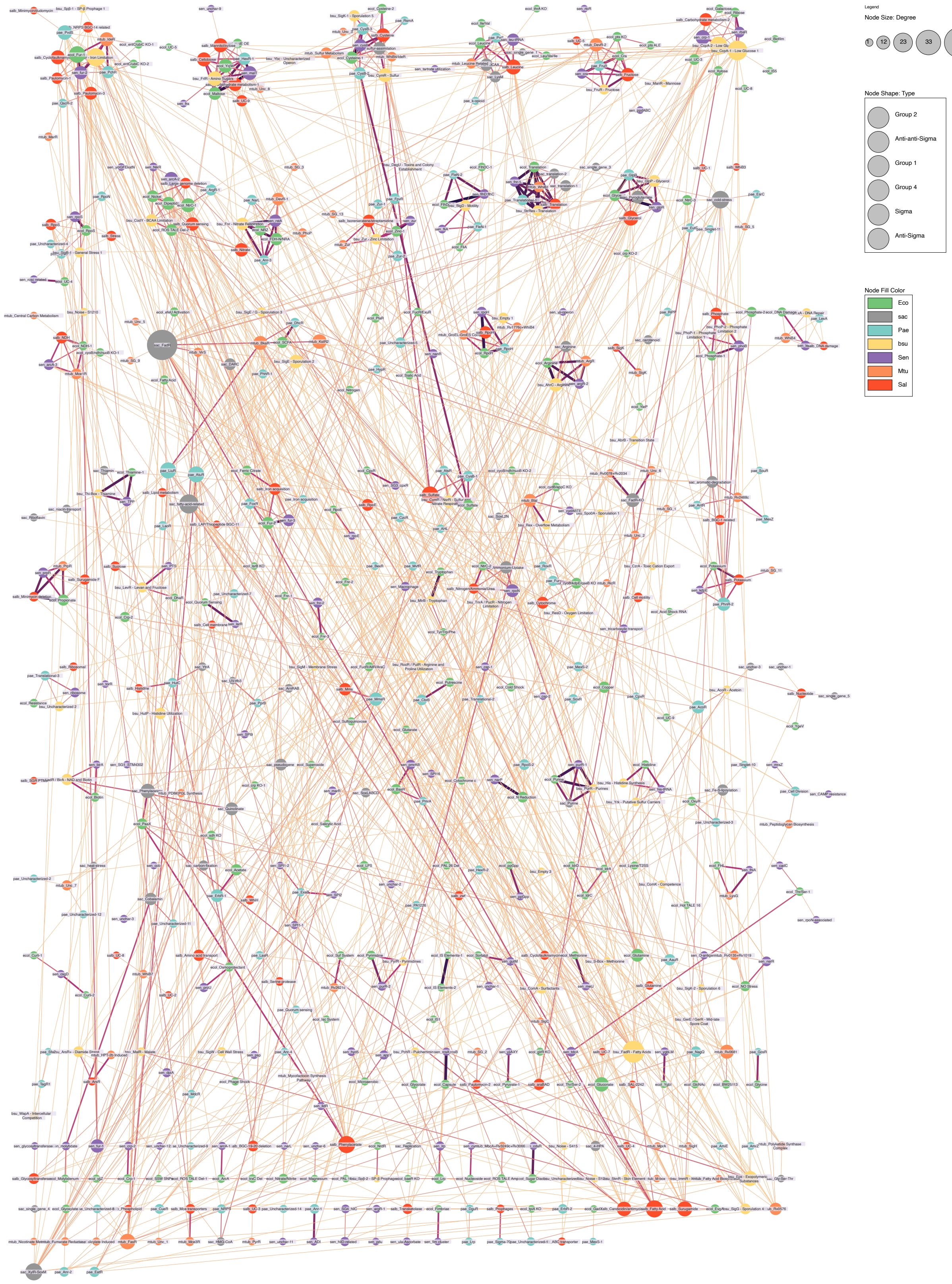


**Supplementary Figure 1**. **Overview of the iModulome.** Each node represents an individual iModulon. Node fill color depicts the species the iModulon was detected in as per the legend. Node size reflects degree and edge thikcness and colour represent edge weight as per legend. The network represents the 1500 most highly weighted edges (< 0.648 edge weight) and was clustered using the MCL clustering method from the clusterMaker app in Cytoscape v.3.10.2.
