## Supplementary figure 2 for "Machine Learning Uncovers the Transcriptional Regulatory Network for the Production Host *Streptomyces albidoflavus*"

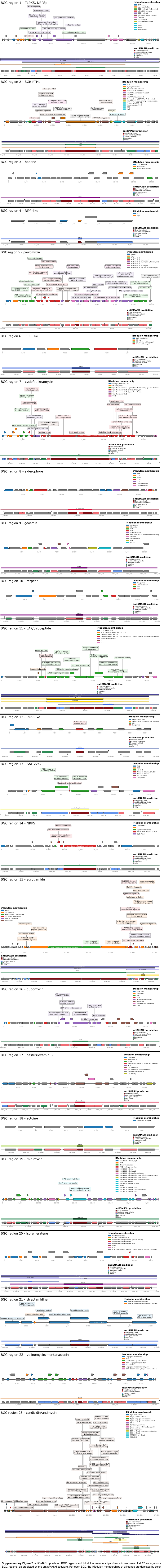


**Supplementary Figure 2.** antiSMASH predicted BGC regions and iModulon memberships. Genomic overview of all 23 endogenous BGC regions, as predicted by the antiSMASH software. For each BGC the iModulon memberships of all genes are depicted by color according to the legend. Genes that are labelled are members of iModulons containing the core biosynthetic genes for respective BGC. The images below are outputs from antiSMASH.
