## Supplementary figure 3 for "Machine Learning Uncovers the Transcriptional Regulatory Network for the Production Host *Streptomyces albidoflavus*"

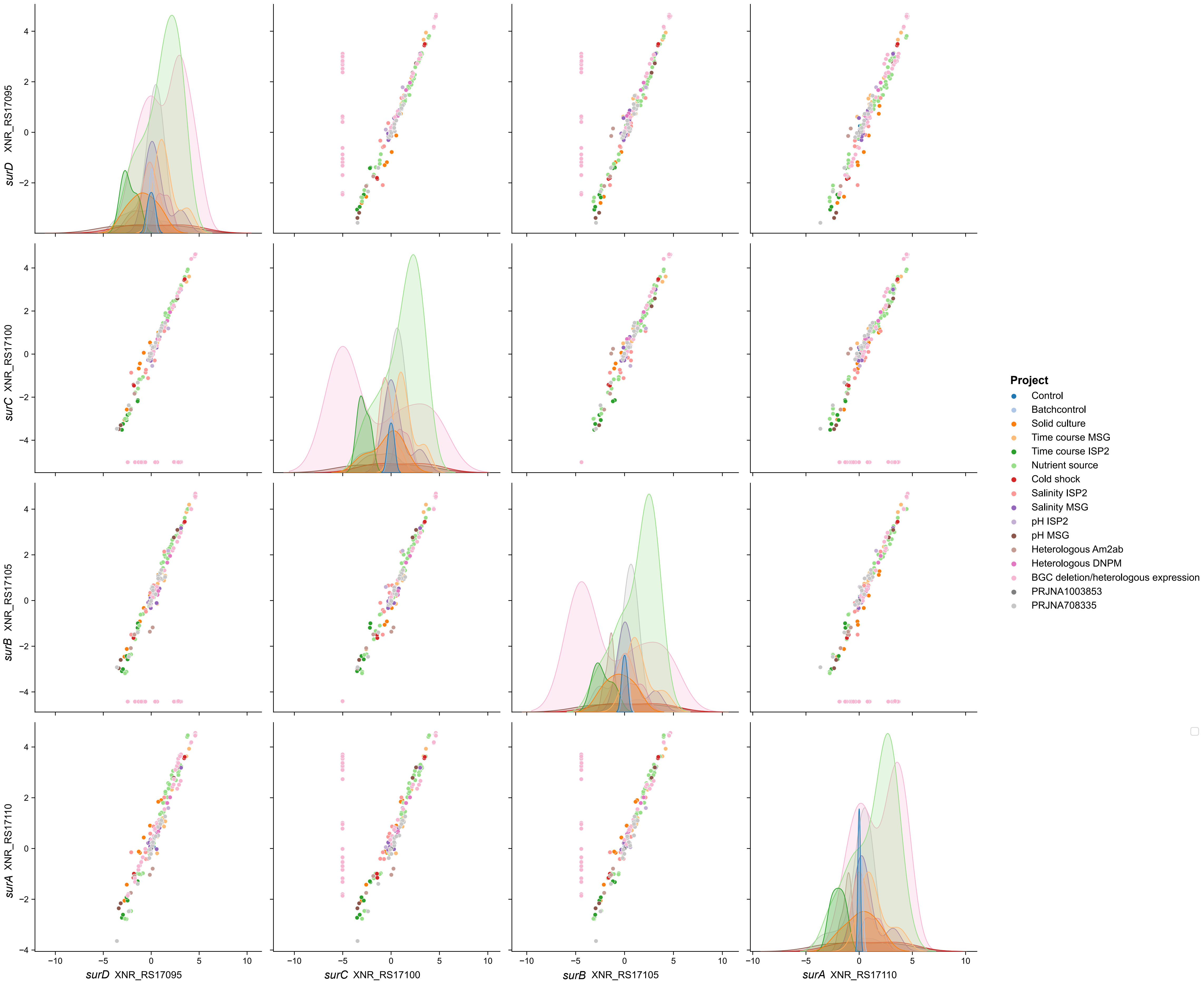


**Supplementary Figure 3. Correlation between *surABCD* activities.** Pairwise plot depicting the gene activities of *surABCD* across different experimental projects (color). The genes in this operon are highly correlated across all conditions except for *surBC* which appear to have been deleted in some of the BGC deletion/heterologous expression samples. This deletion appear to be to main reason why these two genes are not part of the Surugamide iModulon, and instead placed in a separate iModulon (Surugamide F).
